# Dynamic Stiffening to Improve Vasculogenesis of hiPSC-Derived Endothelial Progenitors

**DOI:** 10.1101/2025.10.30.685698

**Authors:** Brett Stern, Bryce Larsen, Katie Halwachs, Kristie Cheng, Davina Tran, Patrik Parker, Nima Momtahan, Aaron Baker, Nicholas Peppas, Adrianne Rosales, Janet Zoldan

## Abstract

Multiple groups have reported on the impact of hydrogel stiffness on vascular network formation *in vitro*, with overall findings indicating that less stiff hydrogels better support vasculogenesis. However, the majority of this research utilized hydrogels with static stiffness, even though vasculogenesis occurs in tandem with changes in extracellular matrix material properties. To that end, we hypothesize that dynamic modulation of hydrogel stiffness during the process of vasculogenesis, recapitulating changes observed during embryonic development, would improve vascular network formation. Using our Collagen I/Norbornene-modified hyaluronic acid hydrogel system, we swelled in additional crosslinker and photoinitiator and exposed the hydrogel to UV light, enabling hydrogel stiffening at user-defined time points with no significant effect on cell viability. We observed that *in situ* stiffening at early time points, prior to the onset of significant cell migration, resulted in more robust vascular network formation relative to unstiffened controls, while stiffening at later time points disrupted existing vascular networks. These trends continued in *in vivo* experiments in nude mice, with cell-laden hydrogels stiffened at early time points resulting in improved blood flow, while those stiffened at later time points had the opposite effect. We hypothesized that this was due to differential impacts of focal adhesion kinase (FAK) activation following in situ stiffening, as supplementation with a Rho kinase inhibitor, downstream of FAK, partially reversed the effects of *in situ* stiffening. Taken together, this research demonstrates the benefits of incorporating dynamic cues into hydrogel design to create more physiologically relevant vasculature.

## Introduction

The vascularization of tissue remains a key challenge in regenerative medicine. As a result, many groups have researched methods required to generate angiogenic biomaterials^1^. One important area of research has been investigating the impact of extracellular matrix properties on vasculogenesis^2^. Focusing on stiffness, multiple groups^3–5^, including us^6–8^, have modulated hydrogel stiffness and came to similar conclusions; an increase in hydrogel stiffness tends to reduce vascular network formation, with vascular network formation not occurring at all at elastic moduli above 3 kPa. This is in contrast to the robust vascular networks that exists *in vivo* in nearly every organ, with stiffnesses ranging from hundreds of pascals to megapascals ^9^.

This discrepancy may be in part because most of the research investigating the impact of matrix stiffness on vasculogenesis was conducted in static systems. During embryonic development, primitive vasculature forms approximately 3 weeks post-fertilization, prior to the development of organotypic ECM^10^, and changes in each organ’s ECM, which includes increases in stiffness, continue throughout embryogenesis as the vasculature continues to develop^11^. This consideration is especially relevant for models utilizing hiPSC-derived endothelial progenitors (hiPSC-EPs), as were used in this study, since hiPSC-EPs show an epigenetic age corresponding to roughly 5 weeks post fertilization^12^. Because of this, creating a system that recreates these dynamic conditions may lead to improvements in vasculogenesis *in vitro*.

To that end, we propose that dynamic modulation of environmental stiffness, as occurs during embryonic development, will improve vasculogenesis. To do this, we utilized our existing Collagen/Norbornene-modified hyaluronic acid interpenetrating polymer network hydrogel (Coll/NorHA IPN)^8^, which allows for stiffness modulation independent of bulk hydrogel density. We then swelled in additional photoinitiator and peptide crosslinker and exposed the hydrogel to UV light to further crosslink it, increasing environmental stiffness. We found that *in situ* stiffening improved vasculogenesis, as measured by our open-source computational analysis pipeline, when performed at early time points and disrupted vasculature when performed at later time points. *In vivo*, we observed similar trends: early-timepoint stiffened hydrogels promoted more extensive vascular networks and also increased blood flow in nude mice. This work demonstrates the utility of dynamic stiffening to improve vessel network formation in 3D systems. Our work emphasizes the need to consider dynamic cues in hydrogel systems to better model the complexity of the endothelial microenvironment.

## Methods

### Maintenance of hiPSCs

hiPSCs (WiCell, DF19-9-11T) were cultured on vitronectin-coated (ThermoFisher Scientific, A14700) six-well plates in Complete Essential 8 medium (E8, A1517001, ThermoFisher Scientific). hiPSCs were passaged upon reaching 70–80% confluency. To passage, hiPSCs were treated with 0.5 mM ethylenediaminetetraacetic acid (EDTA, Invitrogen, 11140-036) for 4.5 minutes at 37°C. EDTA was then removed, and the hiPSCs were resuspended in Complete E8 and seeded as small colonies onto fresh vitronectin-coated plates.

### Generation of a tdTomato-Expressing hiPSC Line

Lentiviral particles containing the sequence p-CAG-tdTomato (Addgene plasmid #83029^13^, a gift from Dr. Stephanie Seidlits) were used to transduce hiPSC. Briefly, 1×10^6^ hiPSCs were transduced with 1 µL of lentiviral stock solution for 16 hours, after which the cells were washed with complete E8 to remove residual virus. 3 days after transduction, tdTomato^+^ hiPSCs were isolated using flow cytometry (MA900 Cell Sorter, Sony) and cultured as described above.

### Diferentiation of hiPSC-EPs

hiPSCs were differentiated into CD34^+^-hiPSC-EPs using a modified protocol from Jalilian *et al*^14,15^. To summarize, hiPSCs were dissociated into a single cell suspension and plated onto Geltrex-coated 6-well plates at a density of 10,000 cells/cm^2^ in Complete E8 supplemented with 10 µM Y-27632. 24 hours after seeding, media was replaced with Complete E8 without Y-27632. 48 hours after seeding, media was replaced with Differentiation Media supplemented with 25 ng/mL Activin A (R&D Systems, 330-AC-050), 30 ng/mL Bone Morphogenic Protein 4 (BMP4, R&D Systems, 314-BP-050), and 0.15 µM 6-bromoindirubin-3-oxime (BIO, Selleckchem, S7198). Differentiation Media consists of Dulbecco’s Modified Eagle Medium (DMEM)/F12 (Cytiva, SH30023.FS), N2 (ThermoFisher Scientific, 17502048), and B27 minus Insulin (ThermoFisher Scientific, A1895601). Differentiation Media supplemented with Activin A, BMP4, and BIO was replaced after 24 hours. 96 hours after seeding, media was replaced with Differentiation Media supplemented with 2 µM SB431542 (Tocris, 1614) and 50 ng/mL Vascular Endothelial Growth Factor (VEGF, R&D Systems, 203-VE-050). Differentiation Media supplemented with these factors was replaced daily for 2 additional days.

### Isolation of CD34^+^-hiPSC-EPs

7 days after seeding, the differentiated cells were incubated with Accutase (STEMCELL Technologies, 07920) for 10 minutes at 37°C and dissociated into single cells. Cells were centrifuged at 300g for 5 minutes and the pellet was resuspended in 200 µL of sorting buffer, Dulbecco’s Phosphate-Buffered Saline (DPBS, GE Healthcare, SH30028.03) containing 2 mM EDTA and 0.5% bovine serum albumin (BSA; Sigma-Aldrich, A8412-100ML), and 2 µL of CD34-PE antibody (Miltenyi Biotec, 130-113-741). Cells were then incubated at 4°C for 10 minutes. Cells were filtered twice with a 35 µm cell strainer (Corning, 352235) to remove cell and extracellular matrix aggregates. CD34^+^-hiPSC-EPs were isolated using a fluorescence-activated cell sorting instrument (S3e; Bio-Rad). Gating and population analysis were performed with Bio-Rad software native to the S3e cell sorter.

### Encapsulation of CD34^+^-hiPSC-EPs in Collagen/NorHA IPN Hydrogels

A total of 500,000 CD34^+^-hiPSC-EPs were centrifuged twice and resuspended in encapsulation media, consisting of Complete Endothelial Growth Media 2 (EGM-2, PromoCell, C-22011) supplemented with 50 ng/mL VEGF, 10 µM Y-27632, and 100 U/mL penicillin-streptomycin (ThermoFisher Scientific, 15140122). Collagen/norbornene-modified hyaluronic acid (NorHA) interpenetrating polymer network hydrogels (IPNs) were synthesized as described previously^8^. In brief, RGD (GCGYGRGDSPG) and an enzymatically degradable (DEG) peptide crosslinker (KCGPQGIWGQCK) were reconstituted in encapsulation media to concentrations of 40 mg/mL and 20 mg/mL, respectively. RGD, DEG, and additional encapsulation media were added to lyophilized NorHA (0.62 norbornene functionalization) to a final concentration of 10 mg/mL NorHA, 2 mM RGD, and either 1.67 mM or 3.34 mM DEG. This DEG concentration crosslinks either 25% or 50% of the available norbornene groups present on the hyaluronic acid, respectively, and we have previously shown that these resulting IPN hydrogels support robust vascular network formation^8^. The photoinitiator lithium phenyl-2,4,6-trimethylbenzoylphosphinate (LAP, Sigma-Aldrich, 900889-1G) was dissolved in encapsulation media to a final concentration of 0.5 wt%. RGD, DEG, and NorHA were synthesized and characterized as described previously^8^.

To encapsulate CD34^+^-hiPSC-EPs, 10x Medium M199 (ThermoFisher Scientific, 11150059), Type I Collagen from Rat Tail (Corning, 354249), LAP, and additional encapsulation media were mixed. The Collagen was then neutralized with 1 M Sodium Hydroxide (Sigma-Aldrich, 72068-100ML). The NorHA solution was then added to the Collagen solution and mixed well, before adding the cell solution to a final concentration of 2.2 million cells/mL. Next, 50 µL of the cell/ IPN pre-hydrogel solution was pipetted into each well of a µ-Slide 18 Well Glass Bottom (Ibidi, 81817) and was allowed to solidify at 37°C at 5% CO_2_. The IPN hydrogels were then exposed to 365 nm UV light at 10 mW/cm^2^ for 50 seconds, and 100 µL of encapsulation media was added. 24 hours after cell encapsulation, media was replaced with EGM-2 supplemented with 50 ng/mL VEGF, termed endothelial culture media, which was replaced daily for 7 days. The final IPN hydrogel contains 1 wt% NorHA and 0.25 wt% Collagen.

### in situ Stifening

To stiffen the cell-laden hydrogels, media was removed and replaced with endothelial culture media supplemented with 3.33 mM DEG and 0.05 wt% LAP for 30 minutes to allow the components to diffuse throughout the hydrogel. At equilibrium, the DEG concentration within the hydrogel will be 1.67 mM, sufficient to crosslink an additional 25% of the total norbornene groups. Media was removed and hydrogels were exposed to 365 nm UV light at 5 mW/cm^2^ for 200 seconds. The hydrogels were then inverted and exposed to UV light under identical conditions. Cell-laden hydrogels were then cultured with EGM2. This process is summarized in **Figure 1**.

**Figure 1:**
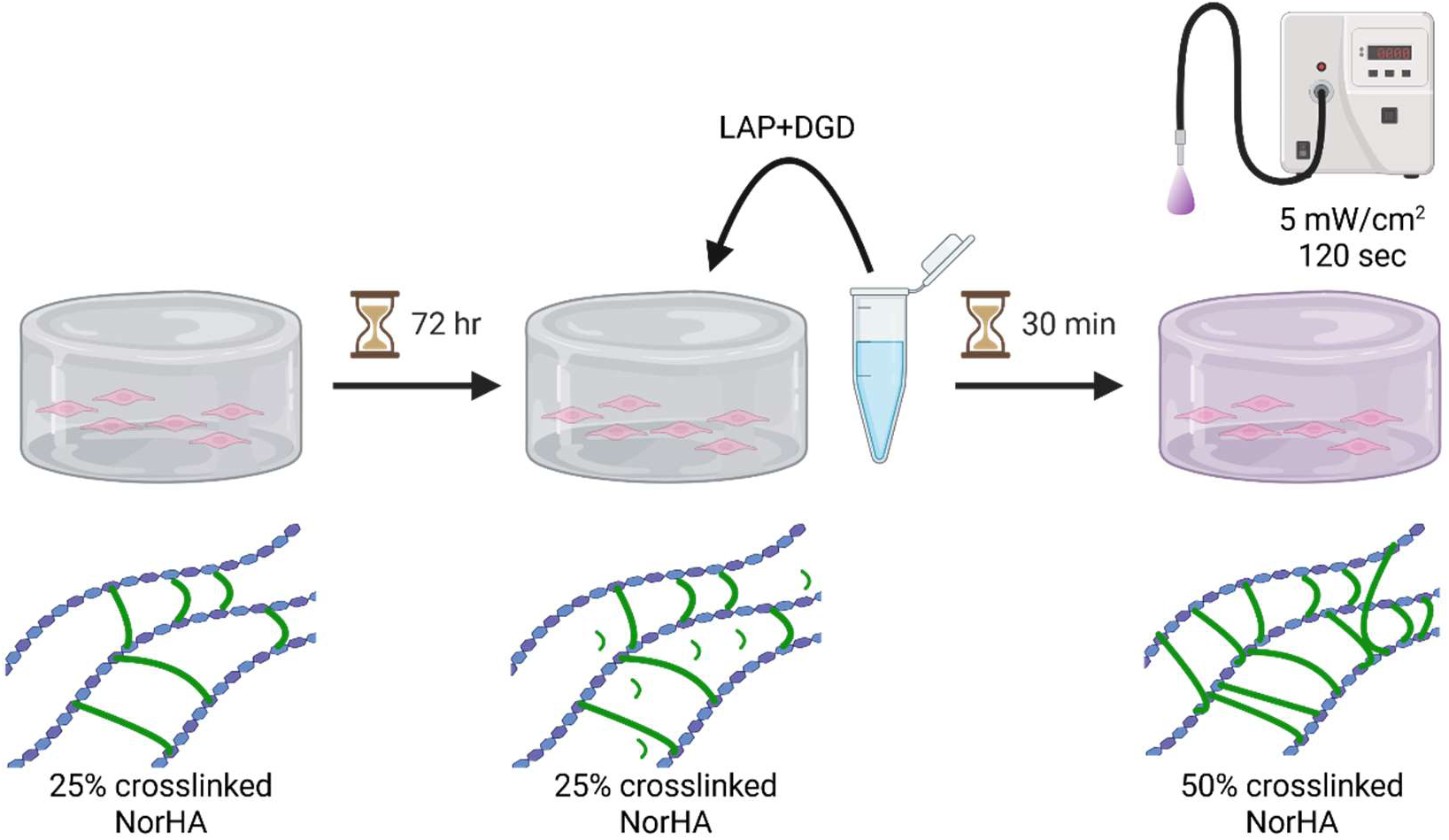
Experimental Overview of the in situ stiffening process. CD34^+^-hiPSC-EPs were encapsulated in 25% crosslinked Collagen/Norbornene-modified hyaluronic acid interpenetrating polymer network hydrogels. At set timepoints after cell encapsulation, additional photoinitiator (LAP) and crosslinker (DEG) were added to culture media and allowed to diffuse throughout the hydrogel for 30 minutes. Following additional UV light exposure, the hydrogel is further crosslinked. Created with BioRender.com.

### Fluorescence Recovery after photobleaching (FRAP)

To determine if *in situ* stiffening occurs evenly throughout the depth of the hydrogel, hydrogel diffusivity was measured at different points in z following a modified protocol by Richbourg *et al*^16^. In brief, 40 kDa Fluorescein-conjugated dextran (ThermoFisher Scientific, D1844) was swelled into Coll/NorHA IPNs for 3 hours at 37°C. Imaging was performed with Zeiss 710 LSM confocal microscope using a 20x objective (NA=0.8, WD=550 µm) with a pinhole size of 1.00 airy units. 3 pre-bleaching images were taken per region of interest, followed by exposing the hydrogel to 100% laser intensity using a 488 nm argon laser. Recovery images were taken at 240 ms per frame with a 0.1-second delay between frames. Images were taken at heights of either 10, 100, or 280 µm from the glass coverslip, representing the bottom, middle, and top of the hydrogels, respectively. 3 images were taken at each height per hydrogel, and each experimental condition consisted of 5 hydrogels. Two-way analysis of variance (ANOVA) was used to evaluate statistical significance.

### Parallel Plate Rheometry

Hydrogel modulus was measured using a Discovery HR-2 rheometer (TA Instruments) with an 8 mm flat stainless-steel geometry and a UV-transparent quartz plate, following an established protocol^8^. Briefly, Coll/NorHA IPNs were generated at initial crosslinking densities of either 25% or 50% in 8 mm diameter acrylic molds and allowed to swell in PBS overnight. A subset of the 25% crosslinked hydrogels were then stiffened as described above, with 6 hydrogels per experimental condition. All modulus measurements were acquired at 1 Hz and 1% strain. One-way ANOVA was used to evaluate statistical significance.

### Live/Dead Staining

Following three days in culture, cell-laden hydrogels were stiffened as described above. The following groups served as controls: hydrogels created at initial crosslinking densities of 25% and 50%, 25% crosslinked hydrogels supplemented with LAP and DEG without UV light exposure, and 25% crosslinked hydrogels exposed to UV light without LAP and DEG supplementation. 24 hours after treatment, hydrogels were stained using 4 µM Calcein AM (Proteintech, PF00008) to visualize live cells and 14.3 nM 4′,6-diamidino-2-phenylindole (DAPI, R&D Systems, 5748/10) to visualize dead cells, by incubation at 37°C for 30 minutes followed by 3 washes with endothelial culture media. DAPI was used instead of the more commonly used ethidium homodimer due to spectral overlap with the cells’ constitutive tdTomato expression. Confocal microscopy was used to image the full depth of the hydrogel, as described previously^8,15,17,18^. Cell viability was quantified using ImageJ. 4 regions of interest were taken per hydrogel, and each experimental condition consisted of 3 hydrogels. One-way ANOVA was used to evaluate statistical significance.

### Daily Monitoring of Vasculogenesis

We performed daily confocal imaging of encapsulated CD34^+^-hiPSC-EPs to determine time points of interest for *in situ* stiffening. CD34^+^-hiPSC-EPs were encapsulated in Coll/NorHA IPNs at crosslinking densities of either 25% or 50% as described above. 24 hours post-encapsulation, we collected confocal z-stacks, with 4 regions of interest per hydrogel and 6 hydrogels per experimental group. This was repeated each day of the 7-day culture period. The ImageJ plugin 3D Suite was used to segment individual cells or cell clusters and measure their compactness to evaluate changes in cell elongation throughout the culture period^19^. One-way ANOVA was used to evaluate statistical significance.

### Network Analysis

CD34^+^-hiPSC-EPs were encapsulated in Coll/NorHA IPNs at 25% crosslinking density and cultured for a total of 7 days. A subset of hydrogels were stiffened either 2, 3, 4, or 5 days after encapsulation. Cell-laden hydrogels created at crosslinking densities of 25% and 50% without *in situ* stiffening served as controls. After 7 days, cell-laden hydrogels were fixed, and confocal z-stacks were collected as described previously by us^8,15,17,18^. Our open-source computational pipeline^18^ was then used to quantitatively evaluate vascular network formation. Two-way ANOVA was used to evaluate statistical significance.

### Reactive Oxygen Species (ROS) Quantification

To measure ROS production during the stiffening process, CD34^+^-hiPSC-EPs were encapsulated in Coll/NorHA IPNs at an initial crosslinking density of 25% and cultured for either 2 or 5 days. On these select days, hydrogels were stiffened to 50% crosslinking as described above. Following stiffening, media was supplemented with 2.5 µM CellRox Deep Red (C10422, Invitrogen) for 30 minutes to visualize intracellular ROS. At this time, a subset of the stiffened hydrogels were also treated with 3 mM of the ROS scavenger n-acetyl cystine (NAC, Sigma-Aldrich, A7250-10G). After 30 minutes, cells were imaged using confocal microscopy. Hydrogels treated with 10 µM hydrogen peroxide (216762, Sigma-Aldrich) for 30 minutes serve as a positive control, and untreated hydrogels serve as a negative control. Intracellular ROS concentrations were quantified using ImageJ by first using the 3D Suite plugin to generate volumes of interest for each cell using their intrinsic tdTomato signal, which was then used as a mask to quantify each cell’s intracellular ROS. One-way ANOVA was used to evaluate statistical significance.

### ROS Scavenging with Network Analysis

To investigate if intracellular ROS negatively affects vascular network formation, we encapsulated CD34^+^-hiPSC-EPs in 25% crosslinked Coll/NorHA IPNs and cultured them for 7 days. *In situ* stiffening was performed either on days 2 or 5, the time points producing the highest and lowest degrees of network formation, respectively, post cell encapsulation. Immediately after stiffening, media was supplemented with 3 mM NAC. The media was replaced with endothelial culture media 24 hours later. Stiffened hydrogels not treated with NAC served as controls. Network analysis was quantified as described above to evaluate if ROS scavenging rescued the cells and allowed for plexus formation. Two-way ANOVA was used to evaluate statistical significance.

### ROCK Inhibition with Network Analysis

To test if ROCK inhibition could disrupt vasculogenesis in hydrogels stiffened at early time points and improve vasculogenesis in those stiffened at later time points, we encapsulated CD34^+^-hiPSC-EPs in 25% or 50% crosslinked Coll/NorHA IPNs and cultured for 7 days. *In situ* stiffening was performed on the 25% crosslinked hydrogels either 2- or 4-days post cell encapsulation. Immediately after stiffening, media was supplemented with 10 µM Y-27632. The media was replaced with endothelial culture media 24 hours later. Hydrogels created at a crosslinking density of 50% served as a control. Network analysis was quantified as described above to evaluate how ROCK inhibition affects vascular network formation. Two-way ANOVA was used to evaluate statistical significance.

### Animal Experiments

CD34^+^-hiPSCs were encapsulated in Coll/NorHA IPNs at initial crosslinking densities of either 25 or 50% and cultured for a total of 7 days. Either 2- or 4-days post cell encapsulation, the 25% crosslinked cell-laden hydrogels were stiffened as described above. On day 7, cell-laden hydrogels were removed from the chamber slides using plastic scoops (VWR, 470125-222) and implanted subcutaneously in mice.

All experimental procedures were approved in accordance with the guidelines set by the University of Texas at Austin Institutional Animal Care and Use Committee.

Four-week-old athymic nude mice (JAX stock #002019) were anesthetized with 2% isoflurane, and four skin incisions were made on the dorsal side of each mouse to create subcutaneous pockets. The pockets were positioned in the top-left, top-right, bottom-left, and bottom-right quadrants with respect to the sagittal and transverse planes. One hydrogel from each of the three experimental conditions (50% crosslinked, stiffened on day 2, stiffened on day 4) was placed into three of the pockets, with the remaining pocket serving as a sham control. Wounds were closed using 4-0 Vicryl resorbable sutures. To establish a baseline for tissue response to the surgical procedure alone, additional mice received incisions and sutures to create subcutaneous pockets, but no hydrogels were implanted. These empty pockets served as a control condition. Perfusion was monitored daily using laser speckle imaging. The hydrogels were harvested, and the animals were euthanized 4 days post hydrogel implantation. A total of 8 mice were used for this experiment.

### Laser Speckle Imaging

Laser speckle contrast imaging was performed to measure blood flow following established protocols^20^. In brief, mice were anesthetized with 2% isoflurane, and the samples were illuminated with a laser diode (50 mW, 785 nm, Thor Labs). Images were captured using a Zoom-7000 lens (Navitar) linked to a Bassler CCD camera (Graftek). Laser speckle images were recorded daily for each mouse. Although five recordings were acquired per session, only one image per day was selected for analysis, corresponding to the frame captured when the animal was not breathing to minimize motion artifacts and ensure accurate measurement of blood flow. The raw speckle data were processed in MATLAB to calculate the relative blood perfusion for each quadrant, normalized to the corresponding sham value and to day 0 (before surgery). Perfusion maps were generated in MATLAB using a JET color scale to maintain consistency across all time points. Statistical analysis was performed using GraphPad Prism 10. A two-way ANOVA was conducted to evaluate the effects of the treatment (column factor) and time (row factor), as well as their interaction, on blood perfusion. Multiple comparisons were analyzed without correction using Fisher’s least significant difference (LSD) test, with a family-wise alpha threshold of 0.05 and a 95% confidence interval.

### Cryosectioning and Imaging of Embedded Hydrogels

Immediately after harvesting, hydrogels were fixed in 4% paraformaldehyde (PFA) and stored at 4°C. PFA was neutralized with the addition of DPBS with 300 mM glycine. Hydrogels were subsequently protected from ice crystal formation by soaking in 10% sucrose in DPBS solution for 24 hours, followed by a 30% sucrose in DPBS solution for 24 hours. The hydrogels were then embedded in optimal cutting temperature compound (OCT) and flash-frozen in isopentane cooled by liquid nitrogen. Hydrogels were sectioned using a cryosectioner (Leica CM1950) at 16 µm thickness and collected on adhesive microscopic slides (Fisherbrand Superfrost Plus). Sections were imaged using confocal microscopy as previously described ^8,15,17,18^. The presence of tdTomato-positive cells confirmed our hydrogel identity, and the tdTomato signal was used to quantify the vasculature structures using the Angiogenesis Analyzer plugin for ImageJ^21^. The Angiogenesis Analyzer was used instead of our custom computational pipeline because the latter is optimized for three-dimensional image stacks, while the cryosections in this study were too thin for 3D reconstruction and required standard two-dimensional angiogenic analysis. Statistical significance was assessed using one-way ANOVA.

### Statistical Analysis

All statistical analysis was performed using GraphPad Prism 10. All analysis was conducted using either one- or two-factor ANOVA followed by a Tukey *post hoc* test for evaluating multiple comparisons. The specific test used is given in relevant sections of the materials and methods. P-values less than 0.05 were considered statistically significant. Significance is denoted as follows: * = p<0.05, ** = p<0.01, *** = p<0.005, and **** = p<0.001. Data are presented as mean ± standard deviation.

## Results

### Transduced hiPSCs and hiPSC-EPs Stably Express tdTomato

Following cell transduction, we isolated tdTomato^+^ cells using FACS. We also confirmed that the transduced hiPSCs expressed both tdTomato and the pluripotency markers Oct4 and Sox2 following extended culture (30+ passages), as shown in **Supplemental Figure 1A-F**, respectively. Successful differentiation into hiPSC-EPs was confirmed by the generation of CD34^+^tdTomato^+^ cells (**Supplemental Figure 1G**) with no impact on differentiation efficiency. These cells were then used for all subsequent experiments.

### FRAP and Rheology Verify Successful in situ Stifening

We first sought to verify whether *in situ* stiffening results in a uniform stiffness throughout the hydrogel depth, given the dependence on UV light penetration through the hydrogel to initiate the stiffening process. To test this, we measured the diffusivity of the hydrogels at different heights using FRAP. Because each condition is identical other than crosslinking density, differences in diffusivity can be correlated with differences in crosslinking. As expected, 25% and 50% crosslinked hydrogels had significant differences in diffusivity at all 3 tested heights (p<0.0001 for 10, 100, and 280 µm), with 50% crosslinked hydrogels having an average of 9.87% lower diffusivity compared to the 25% crosslinked hydrogels, as shown in **Figure 2A**. When hydrogels were exposed to UV light on only the top side of the hydrogel, identical to how UV crosslinking occurs during initial hydrogel formulation, we observed an 11.4% and 9.9% decrease in diffusivity at heights of 100 and 280 µm from the bottom of the hydrogel, respectively (p<0.0001 for both). However, at a height of 10 µm, the diffusivity of the stiffened hydrogel was only 5.7% lower than that of the 25% crosslinked hydrogel (p=0.06). In addition, the diffusivity at 10 µm was 7.93% greater than that at 280 µm (p=0.0067). Taken together, this indicates a stiffness gradient, most likely because of limited UV light penetration to the bottom of the hydrogel. To address this, we stiffened the hydrogels first through UV exposure from the top of the hydrogel to the bottom and then inverted the hydrogel and exposed it to UV light from the bottom to the top. After doing this, we did not observe any difference in diffusivity at any of the three measured heights. At all three heights, the diffusivity was not significantly different from that in the 50% crosslinked hydrogels, indicating that *in situ* stiffening was achieved uniformly, with no detectable stiffness gradient.

**Figure 2:**
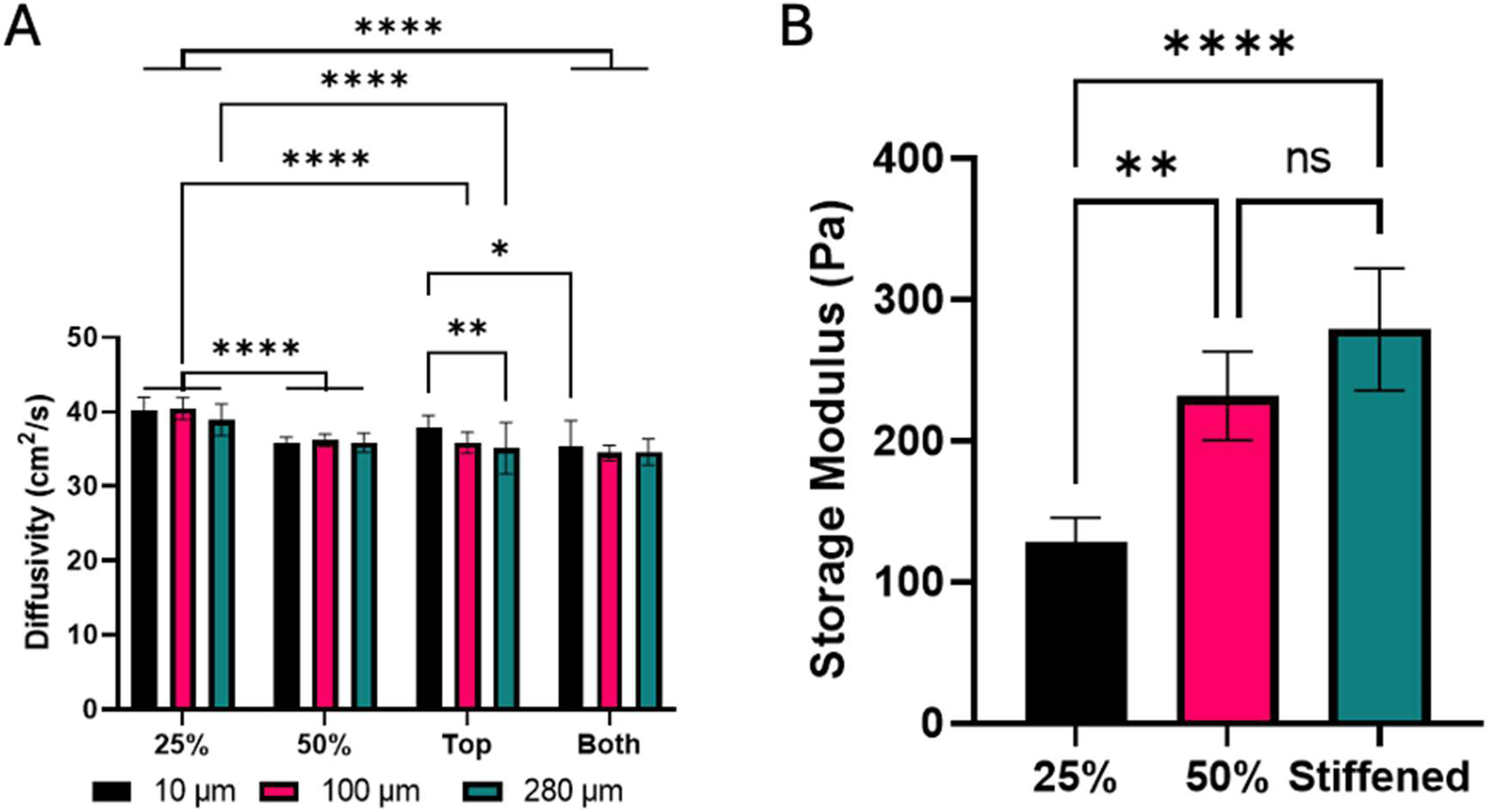
Stifened hydrogels match control in difusivity and storage modulus. (A) FRAP was used to determine if in situ stifening was even throughout the depth of the hydrogel. 25% and 50% crosslinked hydrogels had significantly diferent difusivities at all tested heights. When stifening using the same method used for initial hydrogel formulation (Top) the difusivity at the bottom of the hydrogel was significantly greater than that at the top of the hydrogel, indicating a stifness gradient. The stifness gradient was no longer present when exposing the hydrogel to UV light from both the top and bottom of the hydrogel. n=15 for all conditions. (B) Parallel plate rheometry was used to verify successful stifening. As expected, 50% crosslinked hydrogels had significantly higher stifness than 25% crosslinked hydrogels, and there was no statistically significant diference in storage modulus between the 50% crosslinked hydrogel and stifened hydrogel. n=6 for all conditions.

Next, we measured the stiffness of the *in situ* stiffened hydrogels using parallel plate rheometry, following established protocols^8^, to verify that the *in situ* stiffening not only increased crosslinking density but also achieved the target stiffness. As expected, the 25% crosslinked hydrogels had lower storage moduli than both the 50% crosslinked hydrogels (128.2 Pa vs 231.75 Pa, p=0.0015) and the stiffened hydrogel (128.2 Pa vs 279 Pa, p<0.0001). Most importantly, there was no statistically significant difference in the storage moduli of the 50% crosslinked and stiffened hydrogels (231.75 Pa vs 279 Pa, p=0.13), indicating that the *in situ* stiffening process achieves the desired crosslinking density uniformly throughout the hydrogel and results in a comparable storage modulus as target unstiffened hydrogels (**Figure 2B)**.

### in situ Stifening Is Not Toxic to Encapsulated hiPSC-EPs

We have previously shown that UV crosslinking during initial hydrogel formation results in a decrease in cell viability^8^. Because of this, we were interested in testing whether *in situ* stiffening caused significant cell death. As controls, we included unstiffened hydrogels at both 25% and 50% crosslinking to determine if hydrogel stiffness itself affected cell viability, as well as two conditions that received either LAP and DEG alone or UV light alone as vehicle controls (**Supplemental Figure 2A-E)**. For all tested conditions, cell viability remained high (>90%). In fact, there was no significant difference in cell viability among any of the tested conditions, as shown in **Supplemental Figure 2F**. This demonstrates that *in situ* stiffening does not negatively affect cell viability.

### Encapsulated hiPSC-EPs Reach Maximum Elongation Within 4 Days

During our prior studies encapsulating CD34^+^-hiPSC-EPs in both natural and synthetic hydrogels^8,15^, we observed that vasculogenesis proceeds through a two-step process: first, cells polarize and adopt a spindle shape, and subsequently, they extend and form lumenized, capillary-like structures with adjacent cells. Based on this observation, we aimed to identify which stage of this process is most sensitive to changes in hydrogel stiffness. Specifically, we sought to stiffen the hydrogels at time points corresponding with each of these stages. To determine the appropriate time points for stiffening, we collected confocal z-stacks each day over a 7-day culture period in hydrogels with crosslinking densities of 25% (**Figure 3A-D**) or 50% (**Figure 3F-I**). We then segmented cells in each region of interest and calculated the compactness, a ratio of cell volume to surface area, for each cell using the 3D Suite ImageJ plugin^19^. In the 25% crosslinked condition, we observed an initial 19.6% decrease in compactness between days 1 and 2 (p<0.0001) as cells began to elongate (Figure 3E). Between days 2 and 6, interestingly, there was little change in cell compactness outside of a 3.22% decrease in compactness between days 3 and 4 (p=0.0404). Between days 6 and 7, the 5.55% increase in compactness (p<0.0001) is likely due to issues in segmenting individual cells and instead segmenting clusters of cells. In the 50% crosslinked hydrogels, we observed similar trends, with a 23.2% decrease in compactness between days 1 and 2 (p<0.0001) and a 9.54% decrease between days 2 and 3 (p<0.0001), as shown in **Figure 3J**. After this, there was no significant change in compactness. Overall, these results indicate that cells in both 25% and 50% crosslinked hydrogels rapidly elongate, but this process slows down by days 3-4. Because of this shift from polarization to elongation, we chose to target the 3-4 day range for future experiments. We also tested days 2 and 5 as time points, definitively before and after cell elongation finishes.

**Figure 3:**
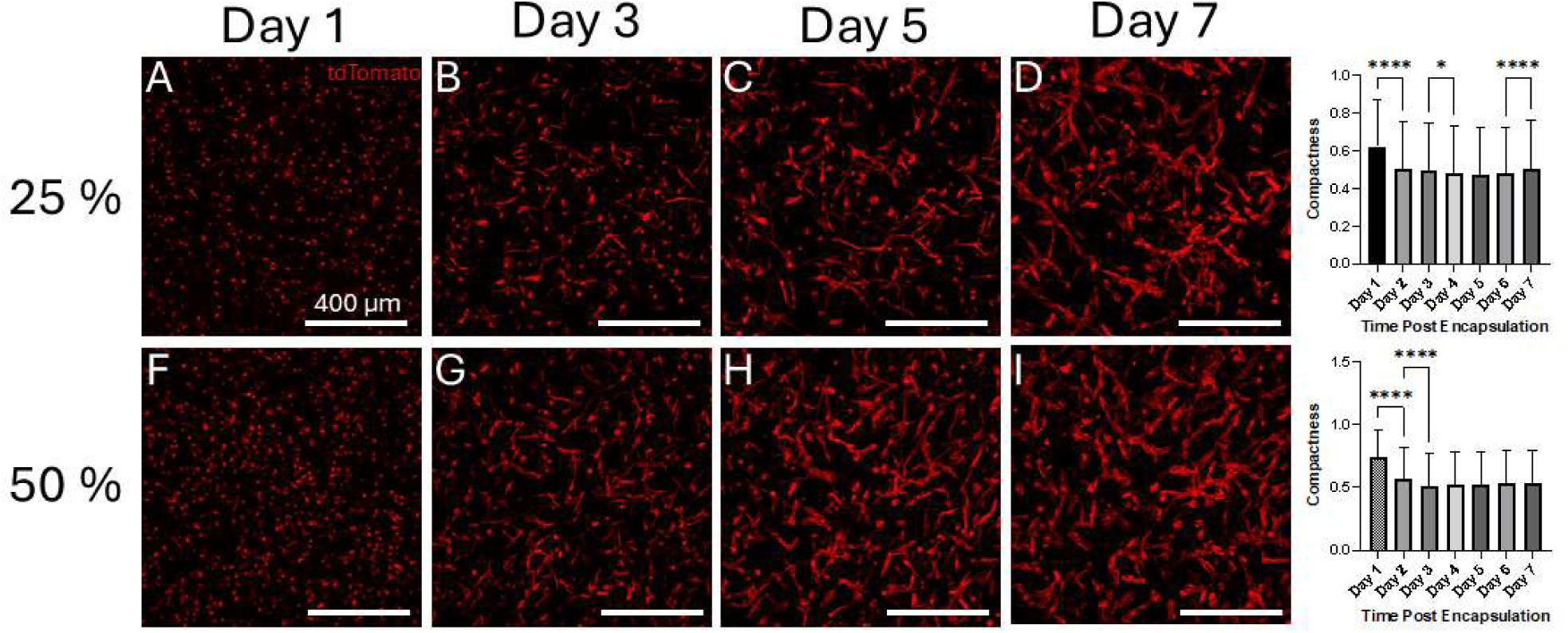
Cell-laden Hydrogels Reach Maximum Elongation by Day 3 or 4 of Culture. CD34^+^-hiPSC-EPs were encapsulated at crosslinking densities of either 25% or 50%. Using the cell’s constitutive tdTomato expression, confocal z-stacks were collected every day for 7 days and ImageJ was used to quantify cell compactness for each day. In both the 25% and 50% crosslinked hydrogels, cell compaction reached a minimum by days 3 or 4. n=24 for all conditions.

### In situ Stifening at Early Timepoints Improves Vasculogenesis

After verifying successful *in situ* stiffening and determining time points of interest for stiffening, we next tested the effects of hydrogel stiffening on vasculogenesis with the following conditions: hydrogels created at initial crosslinking densities of 25% and 50% and hydrogels stiffened either 2-, 3-, 4-, or 5-days post encapsulation (**Figure 4A-E**). We found that, as expected, unstiffened hydrogels at both crosslinking densities formed robust capillary-like vasculature. Interestingly, in the hydrogels stiffened on day 2, we observed a 220% increase in the number of end points, a 237% increase in the number of links between vessels, and a 98.2% increase in the volume fraction of the hydrogel containing vessels relative to the 25% crosslinked control (p<0.0001, p<0.0001 and p=0.0002, respectively). When stiffened on day 3, there was no statistically significant difference in any output from the computational pipeline compared to the 25% crosslinked condition. When stiffened on day 4, we observed a 71% decrease in the number of links and 75% decrease in the volume fraction containing vessels (p=0.0129 and p=0.0079, respectively). When stiffened on day 5, there was a 72% decrease in the volume fraction containing vessels relative to the 25% crosslinked condition (p=0.0116) (**Figure 4F**). Overall, this indicates that *in situ* stiffening can improve vascular network formation when performed at early time points, while disrupting vascular network formation when performed at later time points.

**Figure 4:**
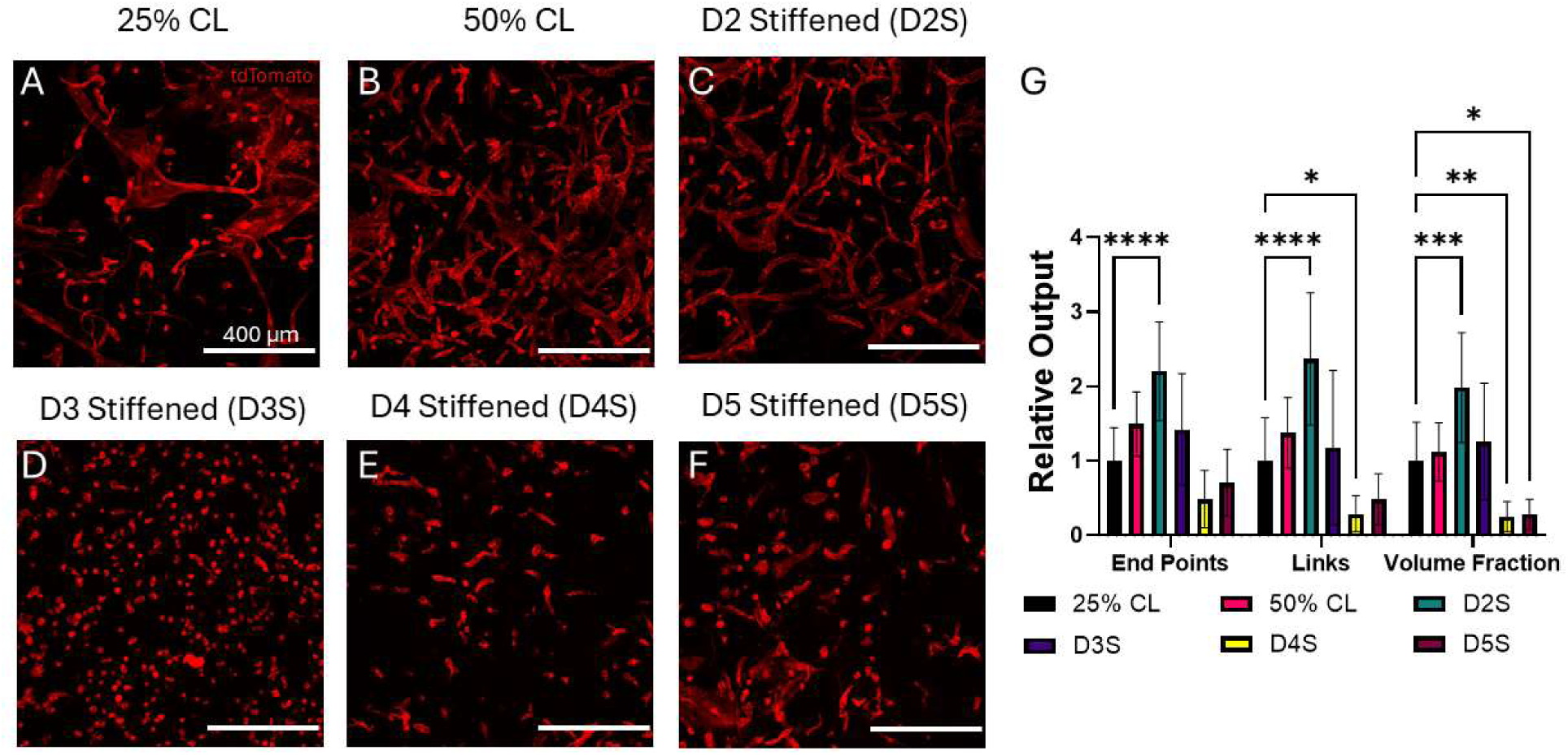
In situ Stifening Improves Vascular Network Formation at Early Time Points. CD34^+^-hiPSC-EPs were encapsulated in Coll/NorHA IPNs at crosslinking densities of either 25% or 50% and cultured for 7 days. A subset of the 25% crosslinked hydrogels was stifened on days 2, 3, 4, or 5 post-encapsulation. Stifening the hydrogels on day 2 increased multiple metrics of vascular network topology, whereas stifening on days 4 or 5, after vasculature had already formed, disrupted vasculature. n=12 for all conditions.

### In situ Stifening Increases Intracellular Reactive Oxygen Species

We hypothesize that reactive oxygen species (ROS) production during the *in situ* stiffening process may be a contributing factor to the reduced vasculogenic potential following stiffening, especially at later time points^22^. This is because increased intracellular ROS activates focal adhesion kinase (FAK) in endothelial cells^23^, and this protein plays an important role in cell migration in part through activation of the Rho/ROCK pathway^24^. We hypothesize that FAK activation early in the process of vasculogenesis would promote cell migration and lead to improved vascular network formation, while FAK activation at later time points would lead to the breakup of existing vasculature. This could therefore explain the differential outcomes of *in situ* stiffening at different time points. To that end, we quantified concentrations of intracellular ROS following *in situ* stiffening at different time points as well as after the addition of the ROS scavenger n-acetyl cystine (NAC). *In situ* stiffening on day 2 of culture resulted in an 82.4% increase in the percentage of ROS^+^ cells (7.36% vs 13.4%, for control and stiffened hydrogels, respectively), although this difference was not statistically significant (p=0.611). Although intracellular ROS remained low following stiffening, supplementation with NAC reduced the percentage of ROS^+^ cells by 98.4% (p<0.0001) (**Supplemental Figure 3**). When stiffening on day 5, there was a 161% increase in intracellular ROS compared to unstiffened cells (p=0.0018), and adding NAC reduced intracellular ROS by 44.7% (p=0.035) to a level similar to that of the control condition (p=0.709) (**Supplemental Figure 4**). We determined that hydrogel stiffening increases intracellular ROS only on day 5, which may play a role in the differential cell response to *in situ* stiffening, and that this increase in ROS can be reduced using a ROS scavenger.

### ROS Scavenging Does Not Afect Vasculogenesis Following in situ Stifening

Given that *in situ* stiffening on day 5 increases intracellular ROS, we next tested whether treatment with n-acetyl cystine could improve vascular network formation. Contrary to our expectations, adding the ROS scavenger had no beneficial effects. NAC supplementation in hydrogels stiffened on day 2 resulted in a 44% decrease in branch points relative to those that did not receive NAC (p=0.015) (**Supplemental Figure 5**). As expected, hydrogels stiffened on day 5 performed significantly worse than those stiffened on day 2, with a 39.5% decrease in the number of end points (p=0.0356), 67.1% decrease in the number of branch points (p<0.0001), and a 46% decrease in the number of links (p=0.01). Stiffening on day 5 with ROS scavenging produced similar results relative to hydrogels stiffened on day 2, with a 49.6% decrease in the number of end points (p=0.0044), 74.1% decrease in the number of branch points (p<0.0001), and a 55.5% decrease in the number of links (p=0.0011). From these results, although increases in intracellular ROS are correlated with reduced vasculogenic potential, they alone do not explain our observed differences in network formation following *in situ* stiffening.

### ROCK Inhibition Reverses Efects of in situ Stifening

Since increased matrix stiffness alone can also activate FAK, signaling through integrin-mediated mechanotransduction^25,26^, we next examined how inhibition of the Rho/ROCK pathway would affect vasculogenesis following *in situ* stiffening. To test this, we treated cell-laden hydrogels stiffened on either day 2 or day 4 with a ROCK inhibitor for 24 hours immediately after *in situ* stiffening. Surprisingly, we found improvements in vascular network formation in cell-laden hydrogels stiffened on day 4 with ROCK inhibitor supplementation, whereas those stiffened on day 2 showed no improvements over unstiffened controls (**Supplemental Figure 6**). Specifically, hydrogels stiffened on day 4 had a 48.9% increase in the number of end points, a 38.6% increase in the number of links compared to unstiffened controls, and a 39% increase in the volume fraction of the hydrogel containing vessels (p=0.0045, p=0.0286, and 0.0269, respectively). However, although we observed increases in three metrics relating to the number of vessel-forming cells, there was a decrease in network connectivity, based on a 2.32-fold reduction in the percent largest network in the day 4 condition relative to the 50% crosslinked control (p=0.0009). Overall, this suggests that the activation of the Rho/ROCK pathway is partially responsible for the differential effects of *in situ* stiffening.

### In situ Stifened Hydrogels Improve Blood Flow in vivo

To investigate whether *in vitro* stiffening can enhance vasculogenesis *in vivo*, we implanted cell-laden hydrogels subcutaneously in nude mice for 4 days. During this period, relative blood flow at each surgical site was quantified using laser speckle imaging. As expected, there were no clear trends in blood flow in the sham condition. Similarly, there was no statistically significant change in blood flow in the 50% crosslinked condition despite this hydrogel supporting robust vascularization *in vitro* (**Figure 5A**). In contrast, hydrogels stiffened on day 2, resulted in a 19.7% increase in blood flow on day 3 (p=0.002) and a 24.8% increase on day 4 (p<0.0001) compared to pre-implantation measurements. However, hydrogels stiffened on day 4 caused a statistically insignificant reduction in blood perfusion during the experiment, with the highest reduction observed on day 2 (24.3% decrease, p=0.4263).

**Figure 5:**
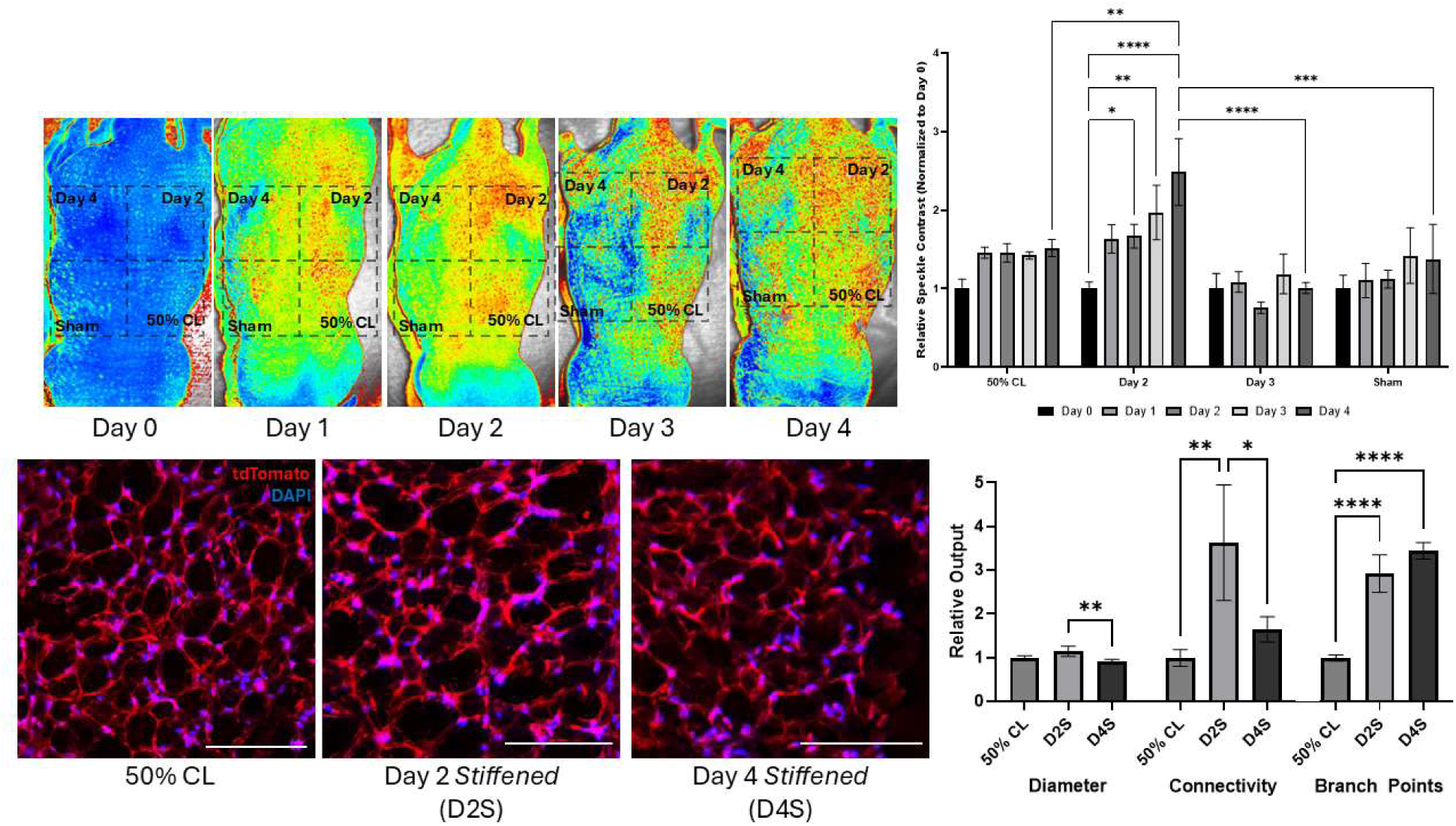
in situ Stifened Hydrogels Improve Blood Flow in vivo. Cell-laden hydrogels, either initially formulated at 50% crosslinking, stiffened on day 2, or stiffened on day 4, were cultured for 7 days under standard conditions. On day 7, hydrogels were implanted subcutaneously in nude mice, and changes in blood flow were quantified using laser speckle imaging. Although 50% crosslinked hydrogels formed robust vasculature in vitro, they provided no beneficial effects in vivo. On the other hand, the day 2 stiffened hydrogels, which form more robust vasculature compared to unstiffened controls, improved relative blood flow on days 3 and 4 post-implantation, whereas hydrogels stiffened on day 4 reduced relative blood flow on days 2 and 3 post-implantation. n=8 for all conditions.

### Pre-implantation stifening recapitulates in vitro vasculogenic trends in vivo

We examined the state of the precultured networks in the various conditions following subcutaneous implantation. Harvested hydrogels were cryosectioned and analyzed for tdTomato presence to confirm the implanted identity of the cells. We then quantified the vascular networks using the Angiogenesis Analyzer plugin for ImageJ (Fig. 5B). In the skeletonized vascular network, points where three or more segments intersected were defined as *branch points*. Connectivity was calculated as the average distance between linear skeleton segments connecting a branch point to an extremity (endpoint). This metric represents the mean distance between successive branch points that are also connected to endpoints, with higher connectivity indicating greater network integration and fewer isolated branches. Network topology in the day 2 stiffened condition showed a modest increase (1.14-fold) in average vessel diameter, a 3.63-fold increase in connectivity, and a 2.93-fold increase in the number of branch points compared to the 50% crosslinked control. The day 4 stiffened condition, conversely, displayed an average diameter 0.90-fold that of the 50% crosslinked control and a non-significant 1.65-fold increase in connectivity. Notably, the day 4 stiffened condition also displayed a 3.45-fold increase in the number of branch points compared to the 50% crosslinked control, indicating a higher prevalence of disconnected clusters. These findings mirror the trends seen *in vitro* and indicate that the day 2 stiffened hydrogels produced a denser, more interconnected vascular structure compared to both the 50% crosslinked control or the day 4 stiffened conditions.

## Discussion

The results of this research demonstrate that dynamic modulation of hydrogel stiffness during the process of vasculogenesis can lead to improvements in vessel network formation and blood flow improvement *in vivo*. Multiple groups, including us^8^, have investigated the effects of extracellular matrix stiffness on vasculogenesis, and the overall findings show that as matrix stiffness increases, vasculogenic potential decreases^2^. However, endothelial cells are found in nearly every organ, meaning that there is functional vasculature in environments with stiffness ranging from hundreds of pascals to millions of pascals. This may be because most research on the subject only investigates the impact of static stiffness, while endothelial cells are exposed to a highly dynamic environment during and after vasculogenesis. To that end, we investigated increasing hydrogel stiffness during the process of vasculogenesis.

*In situ* stiffening has been performed using a range of techniques, including light-induced crosslinking, calcium supplementation for ionic crosslinking, and dynamic covalent bonding^27–30^; these have been used for a variety of applications such as guiding mesenchymal stem cell differentiation and modeling the tumor microenvironment^31^. To our knowledge, only one other group has investigated the impact of *in situ* stiffening on angiogenesis^25^. However, this work was limited in that hydrogel stiffening was only performed after vascular networks had already formed. While post-vascularization stiffening can serve as a model for aging or fibrotic tissue environments, it fails to capture the dynamic mechanical cues that cells experience during the early stages of vasculogenesis. Extracellular matrix stiffness is a critical regulator of endothelial cell behavior, influencing sprouting, migration, and lumen formation^2,6^. By restricting stiffness modulation to after network formation, prior studies may have missed key mechanotransductive events that guide vascular morphogenesis. Therefore, we sought to investigate how temporal modulation of stiffness, specifically during active network formation, affects angiogenesis, with the goal of better recapitulating developmental and regenerative environments where matrix mechanics evolve in parallel with vascular assembly.

One significant limitation of previous studies on *in situ* stiffening is that none, to our knowledge, verified that their changes in stiffness were even throughout the hydrogel. For methods that rely on light-based crosslinking, such as ours, there is a risk of developing a stiffness gradient as less light is able to penetrate thick hydrogels, and methods for characterizing bulk hydrogel material properties, such as rheometry, will not be able to capture this variability. Because of this, we used fluorescence recovery after photobleaching to test the diffusivity of the hydrogel at different points in z, which can then be correlated with hydrogel stiffness. We found that our standard crosslinking protocol failed to increase crosslinking near the bottom of the hydrogel, and we required additional UV light exposure to ensure even crosslinking.

We found that stiffening at an early time point, 2 days post-cell encapsulation, improved vessel network formation, as measured by our vasculogenic computational pipeline, while stiffening had increasingly worse effects when performed at later time points. These effects were also shown *in vivo*, where hydrogels stiffened at early time points increased relative blood flow in mice, while unstiffened hydrogels and hydrogels stiffened at later time points had no effect and negative effects, respectively. Day 2 stiffened hydrogels harvested after implantation in mice also show an increase in both the vessel diameter and connectivity, matching improvements seen *in vitro*. Although we did observe an increase in the number of branch points in the day 4 stiffened condition, this was also accompanied by relatively low connectivity, indicating that the cells formed a splintered plexus less able to support blood flow.

These findings are supported by our knowledge of cell responses during the different stages of vasculogenesis. The cell polarization phase of vasculogenesis is mediated by the extracellular matrix stiffness, because the cells adopt a spindle shape through exerting forces that deform their surrounding ECM^32^. At high stiffnesses, the cells are unable to exert sufficient stress to deform the ECM, resulting in rounded cells that fail to undergo the next step of vasculogenesis^6^. Next, the cells migrate and form lumenized structures with other nearby cells. Contrary to the apical-basal cell polarization stage, higher ECM stiffness tends to promote cell migration because the cells can exert larger magnitude traction forces that enable them to move through their environment^32^. Taken together, *in situ* stiffening on day 2 provides the best of both worlds: a softer environment initially to promote cell extension and a stiffer environment later on to promote cell migration.

We sought to understand why the stiffening at later time points reduced vessel network formation. We hypothesized that this effect was due to the crosslinking reaction generating reactive oxygen species^33^, which are known to be cytotoxic. Although we verified that *in situ* stiffening increased intracellular ROS for hydrogels stiffened on day 5 and that treatment with a ROS scavenger reduced intracellular ROS, we did not observe any improvements in vasculogenesis. We hypothesize that this is because ROS plays an important role in promoting vascular formation rather than being strictly negative, meaning that supplementation with a ROS scavenger may even be detrimental for vasculogenesis^34^.

One alternative explanation for the differential responses to *in situ* stiffening is based on work from other groups demonstrating that increasing matrix stiffness activates focal adhesion kinase (FAK), which has a well-established role in angiogenesis^24,26^. One important effect of FAK activation is phosphorylation of the endothelial tight junction protein VE-Cadherin, resulting in its eventual degradation^24^. This would not have as significant an impact at early points because of a relative lack of cell-cell interactions at this stage. In contrast, it would have an increasingly disruptive effect at later time points as vascular networks formed, in part through VE-Cadherin-containing tight junctions that were more established. One other downstream effector of FAK is RhoA, which is involved in actin polymerization and depolymerization during cell migration^35^. At early time points, RhoA activation would be beneficial for angiogenesis, as it would promote cell migration and would have synergistic effects with increased cell traction forces from the higher ECM stiffness^32^. On the other hand, RhoA activation at later time points would be harmful by promoting cell migration away from already established vasculature, especially when combined with a loss of VE-Cadherin.

To test this, we treated cell-laden hydrogels with a ROCK inhibitor immediately after *in situ* stiffening, either 2- or 4-days post cell encapsulation, and much like our other *in situ* stiffening results, the outcomes were highly time dependent. On day 2, ROCK inhibition appeared to neutralize the beneficial effects of *in situ* stiffening, and none of the outputs of the computational pipeline were significantly different from the unstiffened control. This was expected because other groups have shown that FAK-induced cell migration is necessary for angiogenesis^36^. On day 4, ROCK inhibitor supplementation partially rescued the cell-laden hydrogels from *in situ* stiffening-induced vascular disruption. We observed both a decrease in overall vascular network connectivity and an increase in the number of vessel-forming cells. The decrease in network connectivity is likely because ROCK inhibition does not affect the loss of membrane-associated VE-Cadherin due to FAK activation. The increase in the number of vessel-forming cells may be because ROCK inhibition increases cell proliferation, but more research is needed to confirm this. Taken together, it is likely that these two downstream effects of FAK activation play an important the beneficial or harmful effects of hydrogel *in situ* stiffening.

Overall, this work demonstrates that dynamic modulation of hydrogel stiffness can either improve or disrupt vasculogenesis depending on the stage of vasculogenesis the cells are in, with beneficial effects when the hydrogels are stiffened early and harmful effects when stiffened late. More broadly, this work highlights the importance of incorporating dynamic cues into a hydrogel system to guide vascular network formation.

---

**Supplemental Figure 1:**
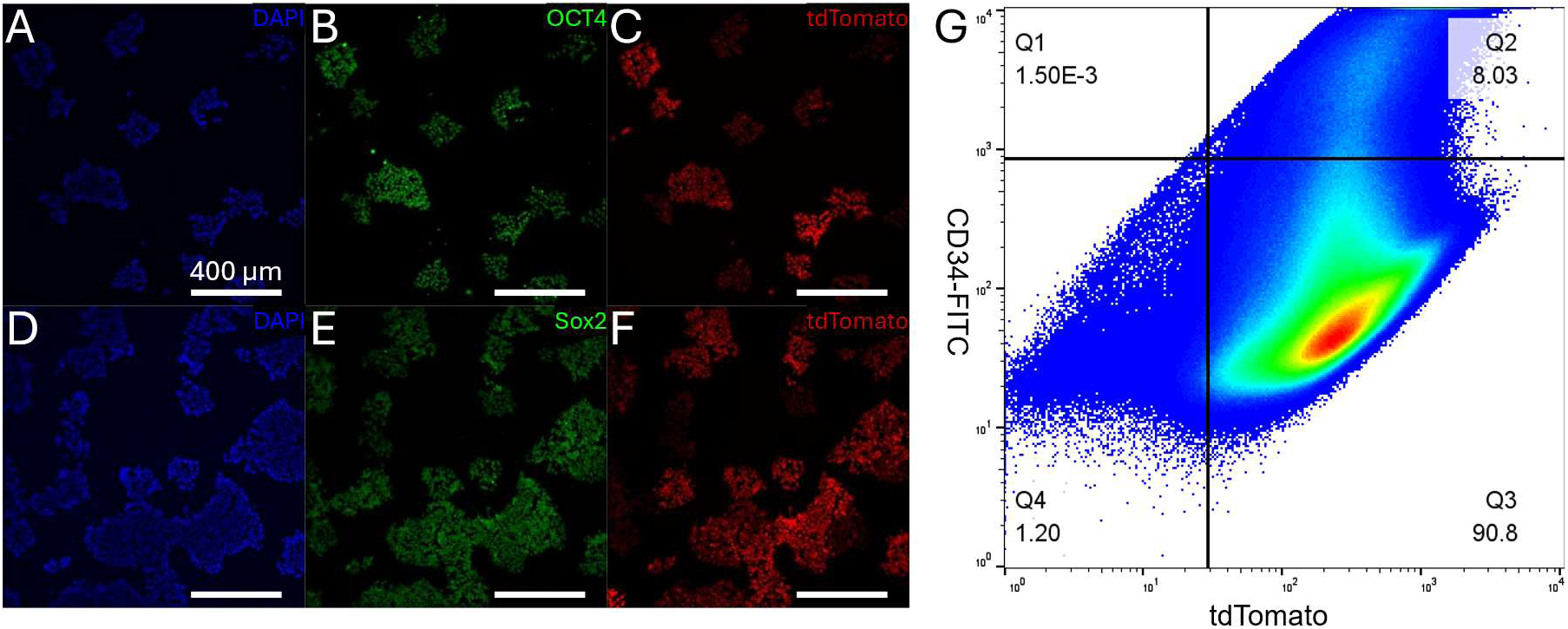
Transduced hiPSCs Stably Express tdTomato. hiPSCs were transduced to express tdTomato using a lentiviral vector. Following extended passage, the cells retain both their tdTomato as well as expression of the pluripotency markers OCT4 (A-C) and Sox2 (D-F). tdTomato expression is also not lost during differentiation into CD34^+^ EPs.

**Supplemental Figure 2:**
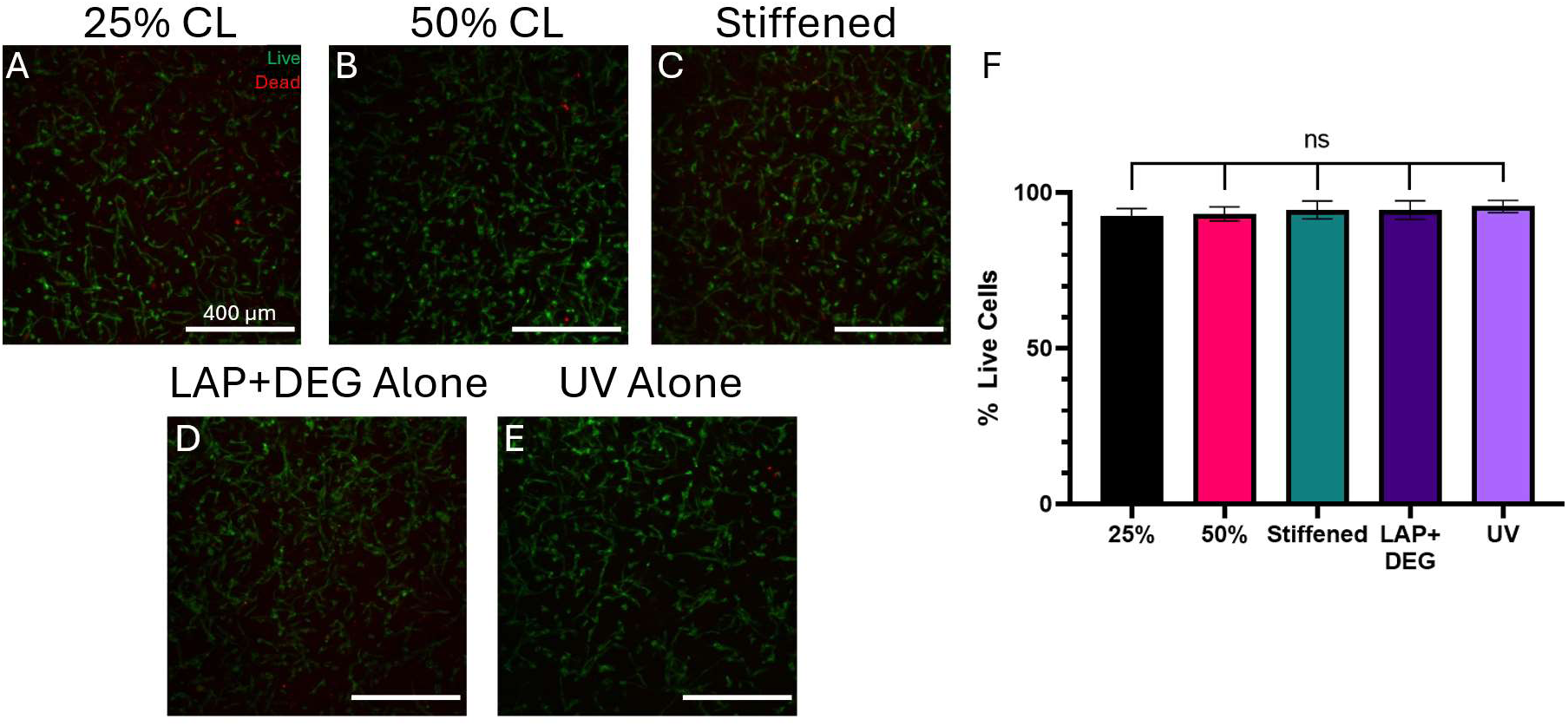
in situ Stiffening is Not Toxic to CD34^+^-hiPSC-EPs. CD34^+^-hiPSC-EPs were encapsulated in Coll/NorHA IPNs at crosslinking densities of either 25% or 50% and cultured for 3 days. A subset of the 25% crosslinked hydrogels were stiffened, and hydrogels treated with either LAP and DEG alone or UV light alone served as vehicle controls. 24 hours after stiffening, LIVE/DEAD staining confirmed no significant differences in viability following in situ stiffening (F). n=12 for all conditions.

**Supplemental Figure 3:**
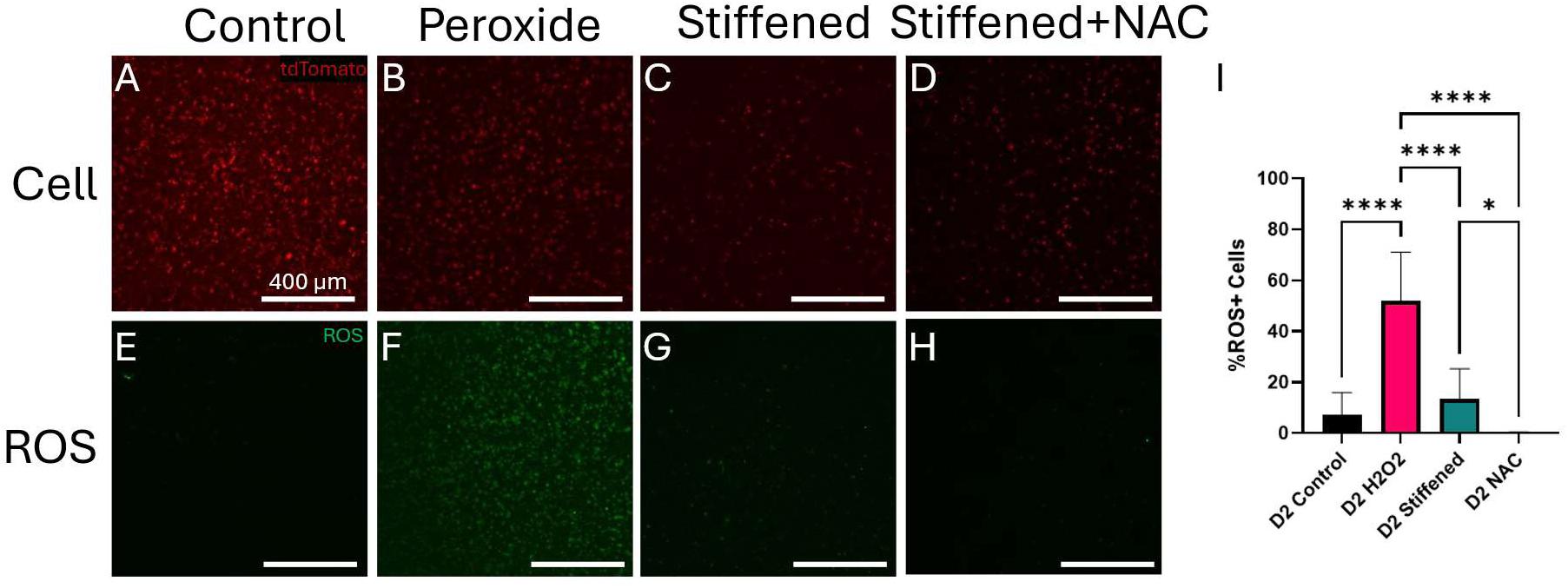
in situ Does not Increase Intracellular ROS on Day 2. Cell-laden hydrogels were cultured for 2 days before receiving either no treatment (A, E), hydrogen peroxide (B, F), in situ stiffening (C, G), or both in situ stiffening and the ROS scavenger NAC (D, H) followed by staining for intracellular ROS. There was no significant change in intracellular ROS following in situ stiffening compared to the control, but addition of NAC was able to reduce intracellular ROS. n=12 for all conditions.

**Supplemental Figure 4:**
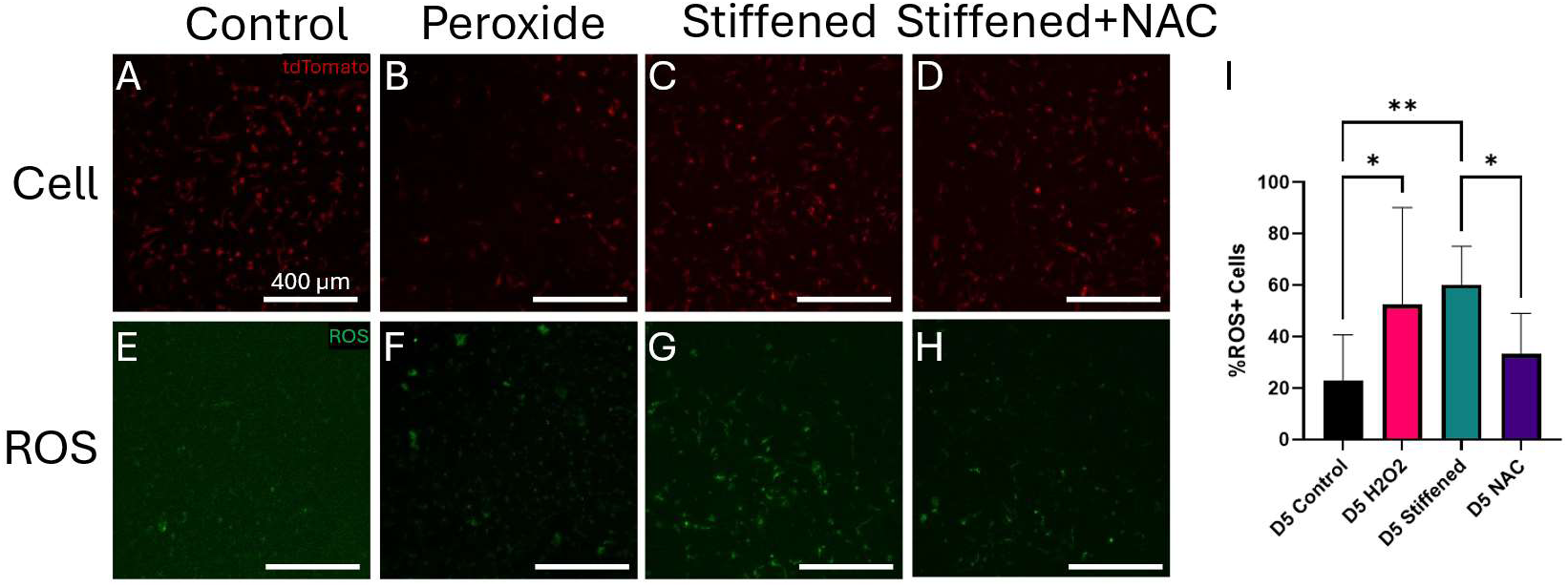
in situ Increases Intracellular ROS on Day 5. Cell-laden hydrogels were cultured for 2 days before receiving either no treatment (A, E), hydrogen peroxide (B, F), in situ stiffening (C, G), or both in situ stiffening and the ROS scavenger NAC (D, H) followed by staining for intracellular ROS. in situ stiffening increased intracellular ROS above levels measured in the hydrogen peroxide positive control, but addition of NAC was able to reduce intracellular ROS. n=12 for all conditions.

**Supplemental Figure 5:**
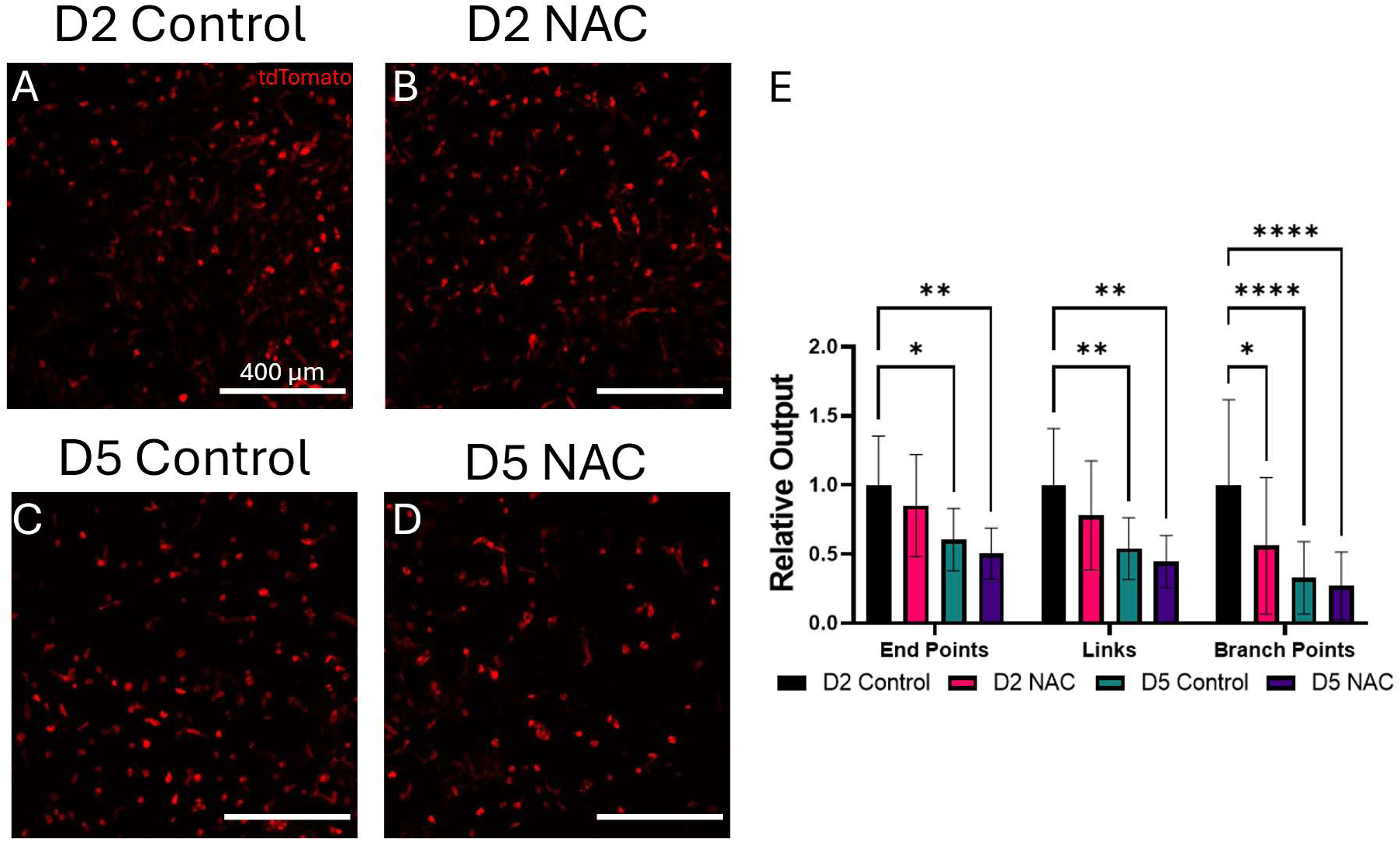
Reducing Intracellular ROS Does Not Improve Vascular Network Formation Following in situ Stiffening. CD34^+^-hiPSC-EPs were encapsulated in Coll/NorHA IPNs at 25% crosslinking and stiffened on either day 2 or 5 post encapsulation. A subset of the hydrogels received NAC on the day of stiffening to determine if reducing intracellular ROS would improve vascular network formation following in situ stiffening. We did not observe any improvements in any metric of our computational pipeline in hydrogels receiving NAC, suggesting that intracellular ROS alone does not play a significant role in vascular disruption following in situ stiffening. n=12 for all conditions.

**Supplemental Figure 6:**
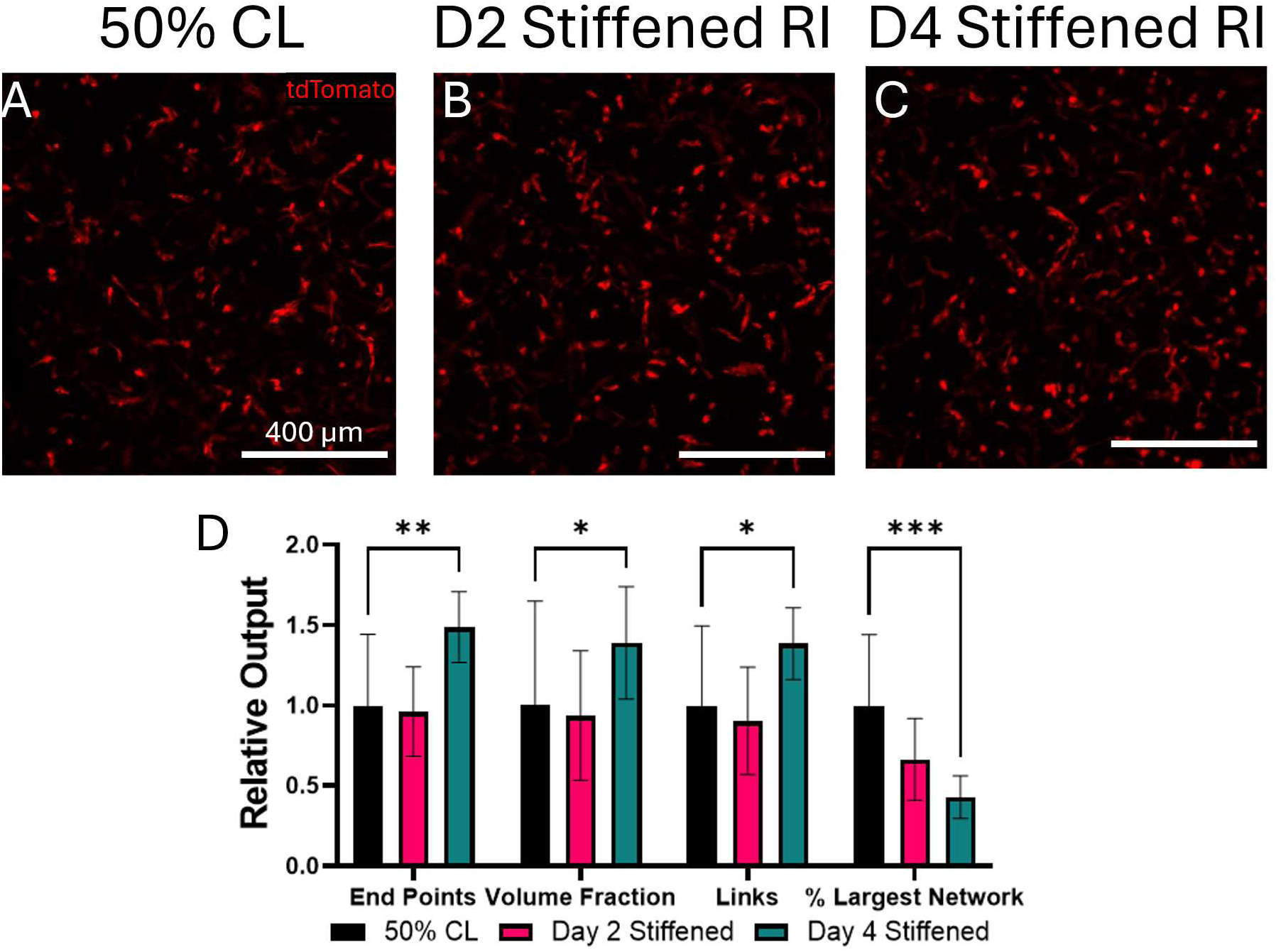
ROCK Inhibition Reverses Effects of in situ Stiffening. CD34^+^-hiPSC-EPs were encapsulated in hydrogels at crosslinking densities of either 25% or 50%. The 25% crosslinked hydrogels were stiffened on either day 2 or 4 and given ROCK inhibitor for 24 hours on the day of stiffening. In the day 2 condition, which normally has more robust vasculature compared to the control, none of the outputs from the computational pipeline were significant different from the 50% crosslinked control, whereas in the day 4 condition, we observed increases in multiple outputs related to the number of vessel-forming cells as well as a decrease in network connectivity. This suggests that ROCK inhibition partially reverses the beneficial effects of early in situ stiffening while rescuing cell-laden hydrogels stiffened at later time points. n=12 for all conditions.

## References

1 Huang, N. F. et al. Bioengineering Cell Therapy for Treatment of Peripheral Artery Disease. Arteriosclerosis, Thrombosis, and Vascular Biology 44, e66–e81 (2024). 10.1161/ATVBAHA.123.318126

2 Crosby, C. O. & Zoldan, J. Mimicking the physical cues of the ECM in angiogenic biomaterials. Regen Biomater 6, 61–73 (2019). 10.1093/rb/rbz003

3 Mason, B. N., Starchenko, A., Williams, R. M., Bonassar, L. J. & Reinhart-King, C. A. Tuning three-dimensional collagen matrix stiffness independently of collagen concentration modulates endothelial cell behavior. Acta Biomaterialia 9, 4635–4644 (2013). 10.1016/j.actbio.2012.08.007

4 LaValley, D. J. & and Reinhart-King, C. A. Matrix stiffening in the formation of blood vessels. Advances in Regenerative Biology 1, 25247 (2014). 10.3402/arb.v1.25247

5 Friend, N. E., McCoy, A. J., Stegemann, J. P. & Putnam, A. J. A combination of matrix stiffness and degradability dictate microvascular network assembly and remodeling in cell-laden poly(ethylene glycol) hydrogels. Biomaterials 295, 122050 (2023). 10.1016/j.biomaterials.2023.122050

6 Han, J. et al. Matrix Stiffness Regulates Mechanotransduction and Vascular Network Formation of hiPSC-Derived Endothelial Progenitors Encapsulated in 3D Hydrogels. bioRxiv, 2025.2004.2011.648340 (2025). 10.1101/2025.04.11.648340

7 Crosby, C. O. K. et al. Quantifying the Vasculogenic Potential of Induced Pluripotent Stem Cell-Derived Endothelial Progenitors in Collagen Hydrogels. Tissue Eng Part A 25, 746–758 (2019). 10.1089/ten.TEA.2018.0274

8 Crosby, C. O. et al. Phototunable interpenetrating polymer network hydrogels to stimulate the vasculogenesis of stem cell-derived endothelial progenitors. Acta Biomater 122, 133–144 (2021). 10.1016/j.actbio.2020.12.041

9 Kretschmer, M., Rüdiger, D. & Zahler, S. Mechanical Aspects of Angiogenesis. Cancers 13, 4987 (2021).

10 Gerecht-Nir, S. et al. Vascular Development in Early Human Embryos and in Teratomas Derived from Human Embryonic Stem Cells1. Biology of Reproduction 71, 2029–2036 (2004). 10.1095/biolreprod.104.031930

11 Abdollahzadeh, F., Khoshdel-Rad, N. & Moghadasali, R. Kidney development and function: ECM cannot be ignored. Differentiation 124, 28–42 (2022). 10.1016/j.diff.2022.02.001

12 Larsen, B., Callahan, C., Rayanki, A., Faulkner, S. & Zoldan, J. Donor Age Impairs Vasculogenic Potential of hiPSC-Derived Endothelial Progenitors. bioRxiv, 2025.2006.2024.661422 (2025). 10.1101/2025.06.24.661422

13 Pathania, M. et al. miR-132 enhances dendritic morphogenesis, spine density, synaptic integration, and survival of newborn olfactory bulb neurons. PLoS One 7, e38174 (2012). 10.1371/journal.pone.0038174

14 Jalilian, E., Raimes, W. & Macias, R. M. Transcriptional profiling reveals fundamental differences in iPS-derived CD34+ Cells versus adult circulating CD34+. Journal of Biology and Medicine 8, 001–013 (2024). 10.17352/jbm.000041

15 Stern, B., Meng, S., Larsen, B., Brock, A. & Zoldan, J. Differentiation Protocol-Dependent Variability in hiPSC-Derived Endothelial Progenitor Functionality. Regenerative Engineering and Translational Medicine (2025). 10.1007/s40883-025-00404-1

16 Richbourg, N. R. & Peppas, N. A. High-Throughput FRAP Analysis of Solute Diffusion in Hydrogels. Macromolecules 54, 10477–10486 (2021). 10.1021/acs.macromol.1c01752

17 Stern, B., Monteleone, P. & Zoldan, J. SARS-CoV-2 spike protein induces endothelial dysfunction in 3D engineered vascular networks. bioRxiv, 2022.2010.2001.510442 (2022). 10.1101/2022.10.01.510442

18 Crosby, C. O. et al. Quantifying the Vasculogenic Potential of Induced Pluripotent Stem Cell-Derived Endothelial Progenitors in Collagen Hydrogels. Tissue Eng Part A 25, 746–758 (2019). 10.1089/ten.TEA.2018.0274

19 Ollion, J., Cochennec, J., Loll, F., Escudé, C. & Boudier, T. TANGO: a generic tool for high-throughput 3D image analysis for studying nuclear organization. Bioinformatics 29, 1840–1841 (2013). 10.1093/bioinformatics/btt276

20 Takematsu, E. et al. Transmembrane stem cell factor protein therapeutics enhance revascularization in ischemia without mast cell activation. Nat Commun 13, 2497 (2022). 10.1038/s41467-022-30103-2

21 Carpentier, G. et al. Angiogenesis Analyzer for ImageJ — A comparative morphometric analysis of “Endothelial Tube Formation Assay” and “Fibrin Bead Assay”. Scientific Reports 10, 11568 (2020). 10.1038/s41598-020-67289-8

22 Yan, R. et al. ROS-Induced Endothelial Dysfunction in the Pathogenesis of Atherosclerosis. Aging and disease 16, 250–268 (2025). 10.14336/ad.2024.0309

23 Usatyuk, P. V. & Natarajan, V. Regulation of reactive oxygen species-induced endothelial cell-cell and cell-matrix contacts by focal adhesion kinase and adherens junction proteins. American Journal of Physiology-Lung Cellular and Molecular Physiology 289, L999–L1010 (2005). 10.1152/ajplung.00211.2005

24 Zhao, X. & Guan, J. L. Focal adhesion kinase and its signaling pathways in cell migration and angiogenesis. Adv Drug Deliv Rev 63, 610–615 (2011). 10.1016/j.addr.2010.11.001

25 Schnellmann, R. et al. Stiffening Matrix Induces Age-Mediated Microvascular Phenotype Through Increased Cell Contractility and Destabilization of Adherens Junctions. Adv Sci (Weinh) 9, e2201483 (2022). 10.1002/advs.202201483

26 Wang, W., Lollis, E. M., Bordeleau, F. & Reinhart-King, C. A. Matrix stiffness regulates vascular integrity through focal adhesion kinase activity. Faseb j 33, 1199–1208 (2019). 10.1096/fj.201800841R

27 Crandell, P. & Stowers, R. Spatial and Temporal Control of 3D Hydrogel Viscoelasticity through Phototuning. ACS Biomaterials Science & Engineering 9, 6860–6869 (2023). 10.1021/acsbiomaterials.3c01099

28 Guvendiren, M. & Burdick, J. A. Stiffening hydrogels to probe short- and long-term cellular responses to dynamic mechanics. Nature Communications 3, 792 (2012). 10.1038/ncomms1792

29 Hull, S. M. et al. 3D bioprinting of dynamic hydrogel bioinks enabled by small molecule modulators. Science Advances 9, eade7880 (2023). doi:10.1126/sciadv.ade7880

30 Stowers, R. S., Allen, S. C. & Suggs, L. J. Dynamic phototuning of 3D hydrogel stiffness. Proceedings of the National Academy of Sciences 112, 1953–1958 (2015). doi:10.1073/pnas.1421897112

31 Zhang, Y. et al. Dynamic Hydrogels with Viscoelasticity and Tunable Stiffness for the Regulation of Cell Behavior and Fate. Materials 16, 5161 (2023).

32 Kretschmer, M., Rüdiger, D. & Zahler, S. Mechanical Aspects of Angiogenesis. Cancers (Basel) 13 (2021). 10.3390/cancers13194987

33 Hoyle, C. E. & Bowman, C. N. Thiol–Ene Click Chemistry. Angewandte Chemie International Edition 49, 1540–1573 (2010). 10.1002/anie.200903924

34 Kim, Y. W. & Byzova, T. V. Oxidative stress in angiogenesis and vascular disease. Blood 123, 625–631 (2014). 10.1182/blood-2013-09-512749

35 Holinstat, M. et al. Suppression of RhoA Activity by Focal Adhesion Kinase-induced Activation of p190RhoGAP: ROLE IN REGULATION OF ENDOTHELIAL PERMEABILITY *. Journal of Biological Chemistry 281, 2296–2305 (2006). 10.1074/jbc.M511248200

36 Tavora, B. et al. Endothelial FAK is required for tumour angiogenesis. EMBO Mol Med 2, 516–528 (2010). 10.1002/emmm.201000106

